# Conditional immobilization for live imaging *C. elegans* using auxin-dependent protein depletion

**DOI:** 10.1101/2021.05.25.445686

**Authors:** Cori K. Cahoon, Diana E. Libuda

## Abstract

The visualization of biological processes using fluorescent proteins and dyes in living organisms has enabled numerous scientific discoveries. The nematode *Caenorhabditis elegans* is a widely used model organism for live imaging studies since the transparent nature of the worm enables imaging of nearly all tissues within a whole, intact animal. While current techniques are optimized to enable the immobilization of hermaphrodite worms for live imaging, many of these approaches fail to successfully restrain the smaller male worms. To enable live imaging of worms of both sexes, we developed a new genetic, conditional immobilization tool that uses the auxin inducible degron (AID) system to immobilize both hermaphrodites and male worms for live imaging. Based on chromosome location, mutant phenotype, and predicted germline consequence, we identified and AID-tagged three candidate genes (*unc-18, unc-104*, and *unc-52*). Strains with these AID-tagged genes were placed on auxin and tested for mobility and germline defects. Among the candidate genes, auxin-mediated depletion of UNC-18 caused significant immobilization of both hermaphrodite and male worms that was also partially reversible upon removal from auxin. Notably, we found that male worms require a higher concentration of auxin for a similar amount of immobilization as hermaphrodites, thereby suggesting a potential sex-specific difference in auxin absorption and/or processing. In both males and hermaphrodites, depletion of UNC-18 did not largely alter fertility, germline progression, nor meiotic recombination. Finally, we demonstrate that this new genetic tool can successfully immobilize both sexes enabling live imaging studies of sexually dimorphic features in *C. elegans*.

**ARTICLE SUMMARY:** *C. elegans* is a powerful model system for visualizing biological processes in live cells. In addition to the challenge of suppressing the worm movement for live imaging, most immobilization techniques only work with hermaphrodites. Here, we describe a new genetic immobilization tool that conditionally immobilizes both worm sexes for live imaging studies. Additionally, we demonstrate that this tool can be used for live imaging the *C. elegans* germline without causing large defects to germline progression or fertility in either sex.

## INTRODUCTION

The discovery of green fluorescent protein (GFP) and subsequent proliferation of both engineered and additional fluorescent proteins has revolutionized biological research by enabling direct visualization of the biological processes occurring within living organisms. This breakthrough changed how scientists viewed cellular processes from snapshots in time generated by fixed images to the complete dynamic picture occurring in real-time within the organism (Chalfie 2009). Live imaging experiments are now performed in nearly every model system spanning all kingdoms of life and can be found in a diverse range of biological fields from molecular biology to systems and synthetic biology (Remington 2011).

For decades, researchers have been curating protocols and technologies to facilitate imaging live *Caenorhabditis elegans* worms. This small, soil-dwelling nematode is completely transparent, thereby making it ideal for live imaging since the intact worm can be placed on a slide and imaged without requiring any tissue dissection or tissue clarification. However, immobilizing the worms on slides can be challenging since wild type worms are highly mobile and spend their life traveling in a sinusoidal pattern across plates eating lawns of bacteria. Further, worms display phototaxis when exposed to light stimulus, avoiding ultraviolet (340nm), blue (470nm), and green (500nm) wavelengths of light (Ward *et al*. 2008). This phototaxis behavior is problematic for live imaging since most of these experiments visualize proteins with GFP, which is typically excited by 488nm light. Thus, this stimulus used to excite GFP also triggers avoidance behavior from the worms.

To enable immobilization of *C. elegans*, custom microfluidic devices have been successfully used to keep worms immobile and alive for imaging experiments anywhere from hours to days (San-Miguel and Lu 2013). While these devices work very well at gently immobilizing the worms, the generation of these devices requires a fabrication facility with the appropriate equipment and expertise in photolithography, which is used to generate the negative molds that create the plastic microfluidic chips. Thus, for many labs without access to in-house fabrication facilities, the generation of microfluidic chips requires external outsourcing, which can be a slow and expensive process if multiple edits of the design are necessary.

Temperature-sensitive hydrogels have also been used effectively to immobilize worms for long timelapse imaging experiments. These hydrogels work by remaining in a viscous liquid state at cold temperature (typically around 4°C) and then solidify at room temperature. This method allows worms to be embedded into the cold hydrogel liquid and become immobilized as the gel warms (Dong *et al*. 2018). Additionally, worms stop thrashing upon cold exposure via ice or cooled liquids (*e*.*g*. a cold hydrogel), which allows for manipulation and placement of the worm within the hydrogel (Chung *et al*. 2008). However, both ice and cold hydrogels can trigger acute cold shock, which is known to trigger a stress response that has negative consequences in worms, including morphological changes within the germline and gut (Robinson and Powell 2016). Therefore, this method is not an option if the worms need to be exposed to elevated temperatures either to induce a conditional mutant or to study the heat shock stress response pathway.

Ultraviolet (UV) crosslinked hydrogels circumvent the need for temperature changes to solidify the gel. Upon exposure to a UV lamp, UV-sensitive hydrogels crosslink together and immobilize worms for long timelaspe imaging (Burnett *et al*. 2018). While this method will not generate any temperature stress responses, exposure to UV causes DNA damage, which can lead to multiple downstream cellular stress events including apoptosis (Mullenders 2018). Thus, UV-sensitive hydrogels may not be an ideal option for studying particular biological processes, such as the maintenance of genome integrity.

The use of agar pads, anesthetics, and polystyrene beads are widely used to immobilize worms. Also, the small size of *C. elegans* adult hermaphrodite, approximately ∼1mm long and ∼0.8mm wide, is about the size of the groove in a LP vinyl record and imprinting these grooves onto an agar pad is effective at immobilizing hermaphrodites when used in combination with anesthetics and polystyrene beads (Rivera Gomez and Schvarzstein 2018). Anesthetics, such as acetylcholine receptor antagonists, have the potential to suppress and/or slow the pharyngeal pumping of the worm, which is analogous to pumping of the mammalian heart. If pharyngeal pumping is suppressed for extended periods of time, then the worm will die. Thus, depending on the specific live imaging experiment, the addition of anesthetics may not be an option. Further, the polystyrene beads, which are usually around 0.1µm and prevent squashing the worm between the coverslip and the agar pad, are small enough for the worm to ingest (Avery and Shtonda 2003; NIKA *et al*. 2016). Currently, it is unclear what the consequences are for the health of a worm when it ingests polystyrene and how this ingestion might affect some biological processes.

In comparison to hermaphrodites, *C. elegans* males are both smaller in length (∼0.8mm) and width (∼0.5mm). Consequently, many methods that immobilize the hermaphrodites fail to immobilize males. *C. elegans* is an excellent model system for sexual dimorphism studies with many sex-specific differences having been identified related to germline processes, nervous system development, and animal aging (Jaramillo-Lambert *et al*. 2007; Barr *et al*. 2018; Hotzi *et al*. 2018; Cahoon and Libuda 2019; Kurhanewicz *et al*. 2020; Li *et al*. 2020). However, due to the differences in body sizes between sexes, it has been difficult to do live imaging-based experiments to analyze and compare the sexually dimorphic features within *C. elegans*. Thus, we developed a genetic immobilization tool that works for both male and hermaphrodite worms. Using the auxin-inducible degron (AID) system, we designed a conditional immobilization system where worms are only paralyzed upon exposure to the plant hormone auxin. Here, we validate this conditional immobilization tool using the *C. elegans* germline and demonstrate that this system works efficiently in both sexes. Notably, we also found that male worms require a higher concentration of auxin to display the same amount of paralysis as hermaphrodites, thereby revealing a potential novel sexual dimorphism within the AID system.

## METHODS

### C. elegans strains, genetics, CRISPR, and culture conditions

All strains were generated from the N2 background and were maintained and crossed at 20**°**C under standard conditions on nematode growth media (NGM) with lawns of *Escherichia coli* (*E. coli*). In Vivo Biosystems tagged *unc-104, unc-18*, and *unc-52* with the auxin inducible degron (AID) tag (amino acid sequence: PKDPAKPPAKAQVVGWPPVRSYRKNVMVSCQKSSGGPEAAAFVK) using CRISPR/Cas9. Both *unc-104* and *unc-18* were tagged on the C-terminus and *unc-52* was tagged on the N-terminus. For all genes, the CRISPR homology-directed repair template was constructed containing 35 base pairs of homology on either side of the insertion site with a small recoded section at the sgRNA site to avoid Cas9 cutting the template. Additionally, a GSTGS amino acid linker was included between the AID tag and each gene. These repair constructs were synthesized as oligos and injected into the worm with two sgRNAs for each gene. All sequences and screening primers for the CRISPR/Cas9 tagging of these genes are in Table S1. All CRISPR/Cas9 worm lines were backcrossed to N2 worms three times before processing with any strain construction.

The following strains were used in this study:

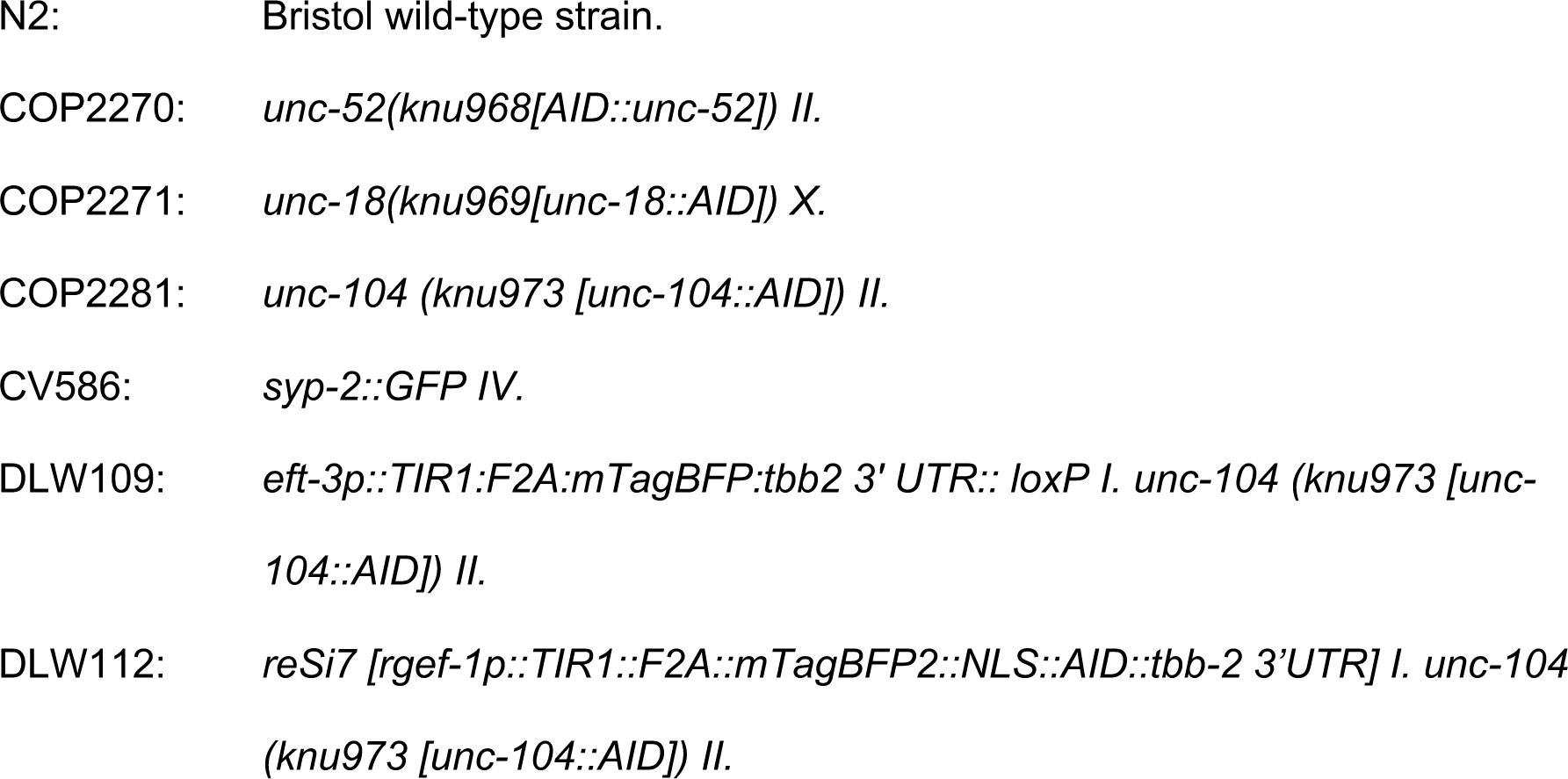
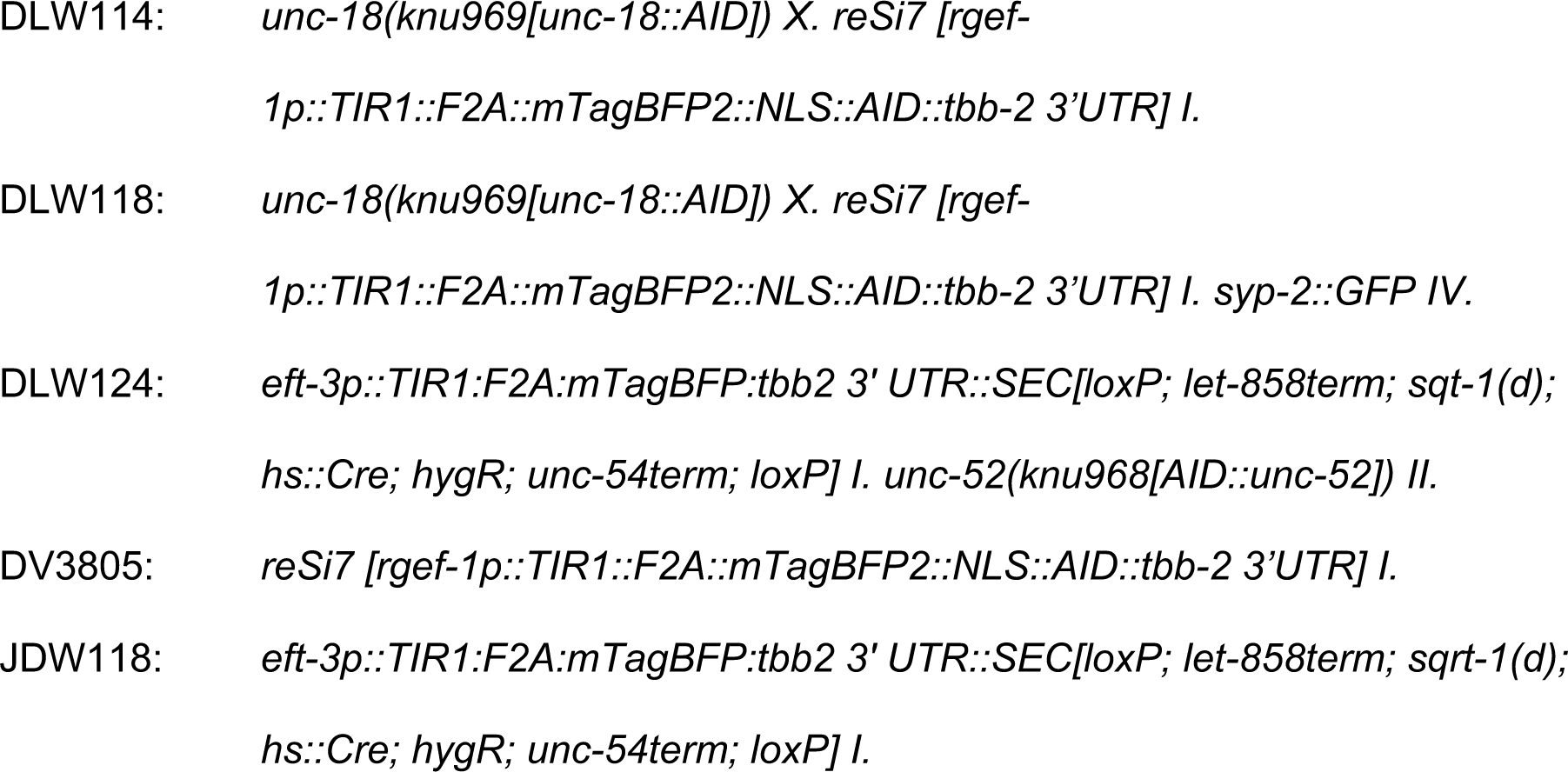

### Worm Tracking

L4 progeny from parental worms on either NGM or NGM with auxin (1mM or 10mM auxin) were transferred to new plates and imaged 18-24 hours later as adults. Worms were imaged using a 1.3-megapixel eyepiece DinoCam camera on a Leica stereoscope with 0.75x magnification. Each movie was captured for 5-10 minutes at 15 frames per second. Using FIJI, movies were background subtracted and a binary image mask of the worms was generated to enable tracking using the FIJI plugin wrMTrck (Nussbaum-Krammer *et al*. 2015). Only worms with long, consecutive durations of tracking were included in the analysis with worms that were tracked for less than 100 frames being excluded from the dataset. Average speed was determined by length the worm traveled in pixels distance divided by the total time the worm was tracked in seconds.

### Fertility assay

To assay fertility, L4 hermaphrodite worms grown on auxin were placed onto new 1mM auxin plates and were transferred every 24 hours for a total of 5 days. After 3 days from removing the hermaphrodite, each plate was scored for living progeny, dead eggs and unfertilized eggs. For each genotype, 12 hermaphrodites were assayed for fertility. If a hermaphrodite went missing, died or bagged during any of the 5 days, then all of her progeny count data was excluded from the dataset. Brood size was calculated for each genotype by summing the living progeny with the dead eggs.

### Conditional immobilization for live imaging

For live imaging experiments, parental worms were placed on plates containing NGM + 1mM auxin or 10mM auxin (Naphthaleneacetic Acid (K-NAA), PhytoTechnology Laboratories, cat no. N610) (Martinez *et al*. 2020). For all male experiments, the parental worms were first picked as L4s onto NGM plates and left to mate for 18-24hrs prior to moving the worms to 10mM auxin NGM plates. L4 progeny grown on auxin were transferred to new auxin plates and young adult progeny worms were used for all imaging experiments. L4 worms from SYP-2::GFP strains were placed at 25°C overnight to enhance GFP expression (Rog and Dernburg 2015; Apttabiraman *et al*. 2017). For mounting of live worms, five to six worms were picked onto poly-lysine coated coverslips (Sigma-Aldrich, cat no. P8920; performed as described in (Wang *et al*. 2018)) (Figure S1). Worms were placed into 1-2µL of imaging media containing M9 media with 25mM serotonin (Sigma-Aldrich, cat no. H7752, (Rog and Dernburg 2015)) and either 1mM auxin (hermaphrodites) or 10mM auxin (males). To prevent the worms from floating in the imaging media, a 7-9% agarose (Invitrogen, cat no. 16500500) pad made from the imaging media was gently placed over the top of the worms. Agarose pads were generated placing a drop of the melted agarose between two coverslips. Then, the solidified pad was lifted with a spatula and gently laid down over the worms. A piece of torn Whatman paper was used to wick excess away the liquid and prevent the agarose pad from floating. A ring of Vaseline was placed around the agarose pad to attach the microscope slide that was placed on top of the coverslip-worm-agarose pad sandwich, and to prevent drying out of the pad while imaging. Then, worms were imaged (see microscopy section). Post-imaging, the worms were monitored for recovery by gently removing the slide from the Vaseline seal with a needle or spatula and either transferring the agarose pad with the worms into a glass well dish filled with M9 media or inverting the pad to pipette M9 media directly on top of the worms. Then, worms were carefully pipetted with low bind tips (Genesee Scientific, cat no. 23-121RS) to NGM plates. Worms were monitored for recovery at 24 hours post imaging. Any worms that did not survive after the recovery were excluded from any analysis. For experiments with worms imaged at 60x, 0.08% tricaine (Ethyl 3-aminobenzoate methanesulfonate; Sigma-Aldrich, cat. no. E10521-50G) and 0.008% tetramisole hydrochloride (Sigma-Aldrich, cat. no. T1512-10G) were added to the M9 imaging media and agarose pads. The addition of the anesthetics did not interfere with the recovery of the worms post imaging.

### Immunohistochemisty

Immunofluorescence was performed as described in (Libuda *et al*. 2013). Briefly, gonads were dissected in egg buffer with 0.1% Tween20 on to VWR superfrost plus slides from 18-24hr post L4 progeny worms from parental worms on NGM only plates or NGM plates with 1mM or 10mM auxin. Dissected gonads were fixed in 5% paraformaldehyde for 5 minutes, flash frozen in liquid nitrogen, and fixed for 1 minute in 100% methanol at −20°C. Slides were washed three times in PBS+0.1% Tween20 (PBST) for 5 minutes each and incubated in block (0.7% bovine serum albumin in PBST) for 1 hour. Primary antibodies (rabbit anti-RAD-51, 1:1500 (Kurhanewicz *et al*. 2020; Toraason *et al*. 2021)) were added and incubated overnight in a humid chamber with a parafilm cover. Slides were then washed three times in PBST for 10 minutes each and incubated with secondary antibodies (goat anti-rabbit AlexaFluor488, ThermoFisher, cat. no. A11034) at 1:200 dilution for 2 hours in a humid chamber with a parafilm cover. Slides were washed two times in PBST then incubated with 2µg/mL DAPI for 15-20 minutes in a humid chamber. Prior to mounting slides were washed once more in PBST for 10 minutes and mounted using Vectashield with a 22×22mm coverslip (no. 1.5). Slides were sealed with nail polish and stored at 4°C until imaged.

EdU staining was performed as described in (Almanzar *et al*. 2021) with minor changes. Briefly, worms were washed three times in PBS + 0.1% TritonX. Then, worms were incubated for 1.5 hours nutating in PBS + 0.1% TritonX with 4mM 5-Ethynyl-2’-deoxyuridine (EdU), which was diluted from a stock 10mM EdU in distilled water from the Invitrogen Click-iT Edu Alexa Fluor 488 imaging kit (Invitrogen, cat. no. C10338). Worms were washed two times in PBS + 0.1% TritonX for 1-2 minutes each then plated onto either NGM or NGM with 1mM or 10mM auxin plates. Time was noted when worms were removed from EdU to start chase time course of the EdU staining. Both male and hermaphrodite worms were dissected and fixed as described above at 0, 10 and 24 hours post removal from EdU and only hermaphrodites were dissected at 48 hours post EdU removal. At each time point 15-20 worms were dissected and washed three times in PBST. Then, slides were either immediately processed with the Click-iT reaction or held in PBST overnight at 4°C and the Click-iT reaction was performed the next day. The Click-iT reaction was performed as described in the kit manual except the volumes in the Click-iT reaction mix were reduced. All slides were incubated with 50µL of the Click-iT reaction mix containing 43µL 1x Click-iT reaction buffer, 2 µL CuSO_4_, 0.2µL AlexaFluor488, and 5µL reaction buffer additive. Slides were incubated for 30 minutes in a humid chamber with a parafilm cover. Then washed three times in PBST for 10 minutes each and incubated with 2µg/mL DAPI in water for 20 minutes with a parafilm cover. Slides were washed once in PBST for 10 minutes then mounted in Vectashield using 22×22mm coverslip (no. 1.5) and sealed using nail polish. All slides were stored at 4°C and imaged within 1-2 days.

### Microscopy

Immunofluorescence slides of gonad stained with RAD-51 were imaged on a GE DeltaVision microscope with a 63x/N.A. 1.42 lens and 1.5x optivar at 1024×1024 pixel dimensions. Images were acquired using 0.2 µm Z-step size and deconvolved with softWoRx deconvolution software. EdU slides and all brightfield timelapses were imaged on GE IN Cell microscope. Edu slides were imaged with a 40x/N.A. 0.95 lens using a Z-step size of 0.72 µm, male brightfield timelapses were imaged with 20x/N.A. 0.75 lens using a Z-step size of 1.3 µm and hermaphrodite brightfield timelapses were imaged 10x/N.A. 0.45 lens using a Z-step size of 3.91 µm. All IN Cell timelapses and images were deconvolved using the IN Cell 3D deconvolution software. *SYP-2::GFP* timelapses were imaged on a Nikon CSU SoRa Spinning Disk Microscope with a 60x water lens/N.A. 1.2 using a Z-step size of 0.3µm.

### Image Analysis and Quantification

RAD-51 gonad images were stitched together using the FIJI (NIH) plugin Stitcher (Preibisch *et al*. 2009) and analyzed in Imaris (Oxford Instruments) as described in (Toraason *et al*. 2021) with minor changes. Only the pachytene region of each gonad was analyzed for RAD-51 foci per nucleus, which was determined by DAPI morphology. The pachytene region was defined by the first row that did not contain more than 1-2 transition zone half-moon like nuclei and the last row that contained all pachytene nuclei with the occasional single diplotene nucleus. These criteria were used for establishing the pachytene region in both hermaphrodites and males. To determine the position of the EdU staining within the germline, the EdU gonad images were max intensity *z*-projected in FIJI. Then, the position of the EdU straining front was determined by the last nucleus within the germline labeled with EdU. Max intensity *z*-projection montages and movies were made in FIJI, and only *GFP::SYP-2* movies were stabilized using the FIJI plugin “StackRegJ_” (https://research.stowers.org/imagejplugins/). This stabilization was necessary to reduce the motion of the germline inside in the worm and generate a stable movie for viewing. Additionally, photobleach correction was applied to the *GFP::SYP-2* male movie using the photobleach correction application in FIJI. All images and movies have been slightly adjusted for brightness and contrast using FIJI.

### Statistics

All statistical tests were performed using Prism. For the worm tracking assay, the average speed of each worm was calculated in the FIJI plugin “wrMTrck” and the multiple comparisons Kruskal-Wallis test was performed to determine statistical differences between each genotype assayed. For the fertility assay, brood size was determined by summing living progeny and dead eggs and statistical differences were determined using 2way ANOVA with Dunnett’s multiple comparisons test. For the RAD-51 and DAPI body quantification, statistical differences were determined using the nonparametric Mann-Whitney test. Each test used is indicated in the Results section next to the reported p-value and all n values are reported in the figure legends.

### Data Availability

All strains are available for request. All supplemental materials are available at Figshare. Figure S1 shows a diagram of the steps used to immobilize worms for live imaging. Table S1 is contains all the primers and sgRNAs used for CRIPSR generations of *unc-18::AID, unc-104::AID* and *AID::unc-52*. Movie S1 is a brightfield timelapse of an immobilized hermaphrodite at 10x magnification. Movie S2 is a timelapse of SYP-2::GFP in an immobilized hermaphrodite at 60x magnification. Movie S3 is an entire germline view of SYP-2::GFP at 60x in an immobilized hermaphrodite. Movie S4 is a brightfield timelapse of an immobilized male at 20x magnification. Movie S5 is a timelapse of SYP-2::GFP in an immobilized male at 60x magnification. Movie S6 is an entire germline view of SYP-2::GFP at 60x in an immobilized male.

## RESULTS

### Reversible paralysis from auxin-dependent depletion of UNC-18 and UNC-104

To conditionally immobilize worms, we used the auxin-inducible degron (AID) system, which has been used in multiple different worm tissues to selectively deplete proteins of interests at specific stages of development (Zhang *et al*. 2015; Pelisch *et al*. 2017; Kasimatis *et al*. 2018; Serrano-Saiz *et al*. 2018; Martinez *et al*. 2020; Ashley *et al*. 2021). This system requires three components: (1) a protein containing the AID sequence; (2) expression of the plant F box protein Transport Inhibitor Response 1 (TIR1), which can be regulated using tissue-specific promotors; and (3) the plant hormone auxin, which can be absorbed externally by the worms (Figure 1A). Auxin exposure promotes the binding of TIR1 to the degron sequence. TIR1 is able to interact with components of the endogenous SCF E3 ubiquitin ligase complex generating a functional complex that can ubiquitinate the degron tagged protein and target it for proteasome-mediated degradation (Natsume and Kanemaki 2017). Additionally, this protein degradation is completely reversible once the worms are removed from auxin, such that after a period of time the degron tagged protein can return to wild type levels (Zhang *et al*. 2015).

**Figure 1.**
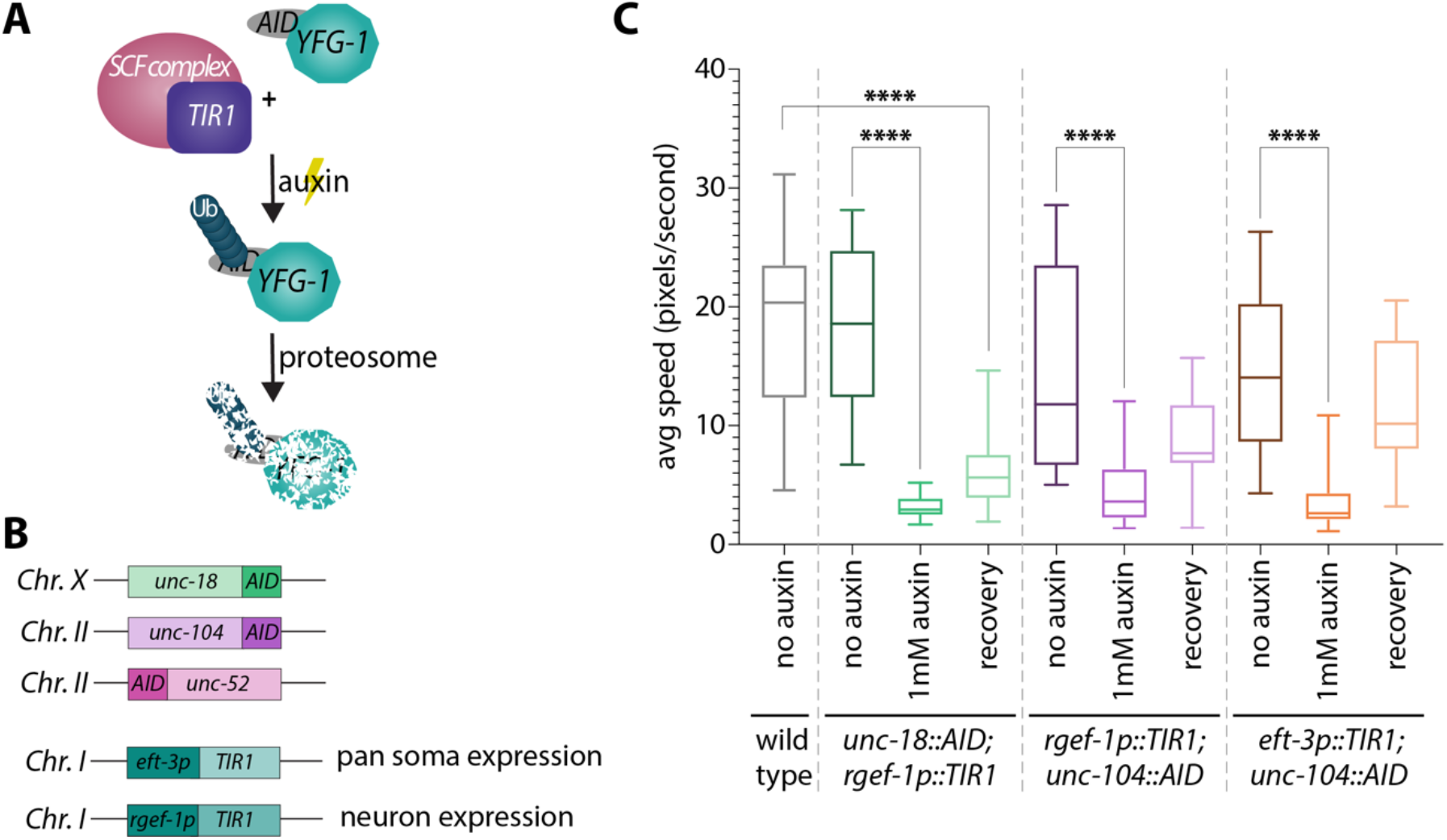
Auxin-dependent depletion of *unc-18* and *unc-104* permits conditional immobilization of hermaphrodites. **(A)** Diagram of the auxin inducible degron (AID) system showing how *TIR1* associates with the endogenous SCF E3 ligase complex that in the presence of auxin cause ubiquination (Ub) of the AID::YFG-1 protein (Your Favorite Gene 1). This ubiquination results in proteosomal degradation of AID::YFG-1. **(B)** Schematics of the three candidate genes CRISPR/Cas9 tagged with AID and the *TIR1* constructs used with the expression location of *TIR1* based on the *eft-3* or *rgef-1* promotors. The chromosome number where each construct is located in the genome is indicated on the left of each schematic. **(C)** Quantification of the average speed in pixels per second of wild type (n= 30), *unc-18::AID; rgef-1p::TIR1* (no auxin n=18, 1mM auxin n=16, recovery n=26), *rgef-1p::TIR1; unc-104::AID* (no auxin n=24, 1mM auxin n=31, recovery n=18), and *eft-3p::TIR1; unc-104::AID* (no auxin n=15, 1mM auxin n=26, recovery n=24) on plates containing no auxin and 1mM auxin. The recovery category indicates worms that have been off auxin for 18-24 hours prior to tracking worm motion. **** indicates P<0.00001, Kruskal-Wallis.

To conditionally immobilize worms, we combined the AID system with genes that cause severe worm paralysis when mutated and applied this immobilization to visualize the germline in live animals. To narrow down the candidate list of genes, we focused on Chromosomes *X* and *II* since we wanted to use this system to study the *C. elegans* germline and these chromosomes are mainly devoid of germline expressed genes (Reinke *et al*. 2000). We then obtained mutants from all the identified genes on these chromosomes that were indicated on the CGC as being homozygous viable and severely paralyzed. Additionally, we avoided any genes that had the potential to alter the germline, vulval development, or vulval function. From these candidates, we selected three genes to tag with the AID sequence: *unc-104, unc-18*, and *unc-52* (Figure 1B). UNC-104 is a kinesin-3 family motor protein that is primarily involved in transporting synaptic vesicle precursors within neurons (Hayashi *et al*. 2019). UNC-18 belongs to the Sec1p/Munc18 family proteins and plays a critical role in synaptic exocytosis in neurons (Park *et al*. 2017). UNC-52 is an extracellular matrix heparan sulphate proteoglycan that plays an essential role in myofilament assembly of body-wall muscles (Rogalski *et al*. 2001).

Using CRISPR/Cas9, each of these genes were tagged with the AID sequence and genetic crosses were performed to incorporate *TIR1*. Two different *TIR1* constructs were used in this study: 1) *rgef-1p::TIR1*, which expresses only in neurons; and, 2) *eft-3p::TIR1*, which has pan-somatic expression. For all experiments, we found that animals needed to be grown for a single generation on plates containing nematode growth media (NGM) with auxin to exhibit the strongest paralysis phenotype (see Methods).

We first assayed *unc-104::AID* and *unc-18:AID* using the neuron-specific expression of *TIR1 (rgef-1p::TIR1)* and *AID::unc-52* with pan-somatic expression of *TIR1 (eft-3p::TIR1)* (Figure 1B). *AID::unc-52* displayed no changes in mobility when grown on auxin plates, thus this gene was excluded from any further studies. Both *unc-104::AID* and *unc-18::AID* display significant decreases in mobility on auxin plates, which we assayed by tracking the motion of the worms on normal NGM plates and NGM with 1mM auxin (Figure 1C, P<0.0001, Kruskal-Wallis). We noticed that *unc-104::AID* did not display as strong of a mobility defect as *unc-18::AID* when depleted using the neuron specific *TIR1* (median average speed: 3.604 and 2.927 pixels/second, respectively). Moreover, depleting UNC-104::AID with the pan-somatic *eft-3* driven TIR1 exhibited a similar degree of mobility defects to neuron-specific depletion of *unc-104::AID* with the *rgef-1* driven TIR1 (P>0.999, Kruskal-Wallis multiple comparisons test). Although the effects of each driver on UNC-104::AID depletion were statistically indistinguishable, the median average speed was slightly lower with the pan-somatic *eft-3* driver (neuron-specific *rgef-1* driven TIR1: 3.604 pixels/second; pan-somatic *eft-3* driven TIR1: 2.625 pixels/second).

With this worm immobilization technique, worms can be recovered post-imaging and assayed for viability off auxin. We found that upon removal from auxin for 18-24 hours both *unc-104::AID* and *unc-18::AID* worms recovered some degree of normal motion (Figure 1C). *unc-104::AID* with *TIR1* driven by both the neuron-specific *rgef-1* promoter or the pan-somatic *eft-3* promoter recovered motion to levels statistically indistinguishable from wild type, but the overall median average speed in these animals was lower than wild type worms (wild type: 20.34 pixel/second; *rgef-1p:TIR1; unc-104::AID*: 7.669 pixel/second; *eft-3p::TIR1; unc-104::AID*: 10.16 pixel/second). *unc-18::AID* worms were able to recover some motion off auxin compared to the *unc-18::AID* in the presence of auxin; however, this recovered motion was significantly slower than wild type worms (median average speed for *unc-18::AID* on auxin: 2.927 pixel/second; *unc-18::AID* recovery: 5.634 pixel/second; wild type: 20.34 pixel/second; P<0.0001, Kruskal-Wallis). Taken together, these motion recovery experiments suggest that *unc-18::AID* may require more time to completely recover wild type motion compared to the *unc-104::AID* worms. Overall, both *unc-104::AID* and *unc-18::AID* are able to both partially recover movement after removal from the auxin treatment.

### UNC-104::AID depletion has slight behavioral and fertility defects

To examine the effectiveness of this conditional immobilization system for live imaging, we focused on implementing this system for live imaging the *C. elegans* germline. Since the effects of UNC-104 or UNC-18 loss on germline function are unknown, we examined multiple aspects of germline biology to determine if a loss of UNC-104 or UNC-18 causes germline-specific defects. We began by assaying the fertility of hermaphrodite worms containing *unc-104::AID* and *unc-18::AID* under both depletion (in the presence of auxin) and wildtype conditions (Figure 2). For these fertility assays, we counted the number of living progeny, dead eggs and unfertilized eggs from hermaphrodite worms that were moved each day for five days to new plates (see Methods). Scoring fertility over multiple days allowed for observation of most of the hermaphrodite reproductive lifespan, which begins with a large abundance of living progeny and subsequently ends with unfertilized eggs once the hermaphrodite sperm is depleted (Ward and Carrel 1979). To compare the fertility of each genotype, we calculated brood size of each worm, which is the sum of living progeny and dead eggs.

**Figure 2.**
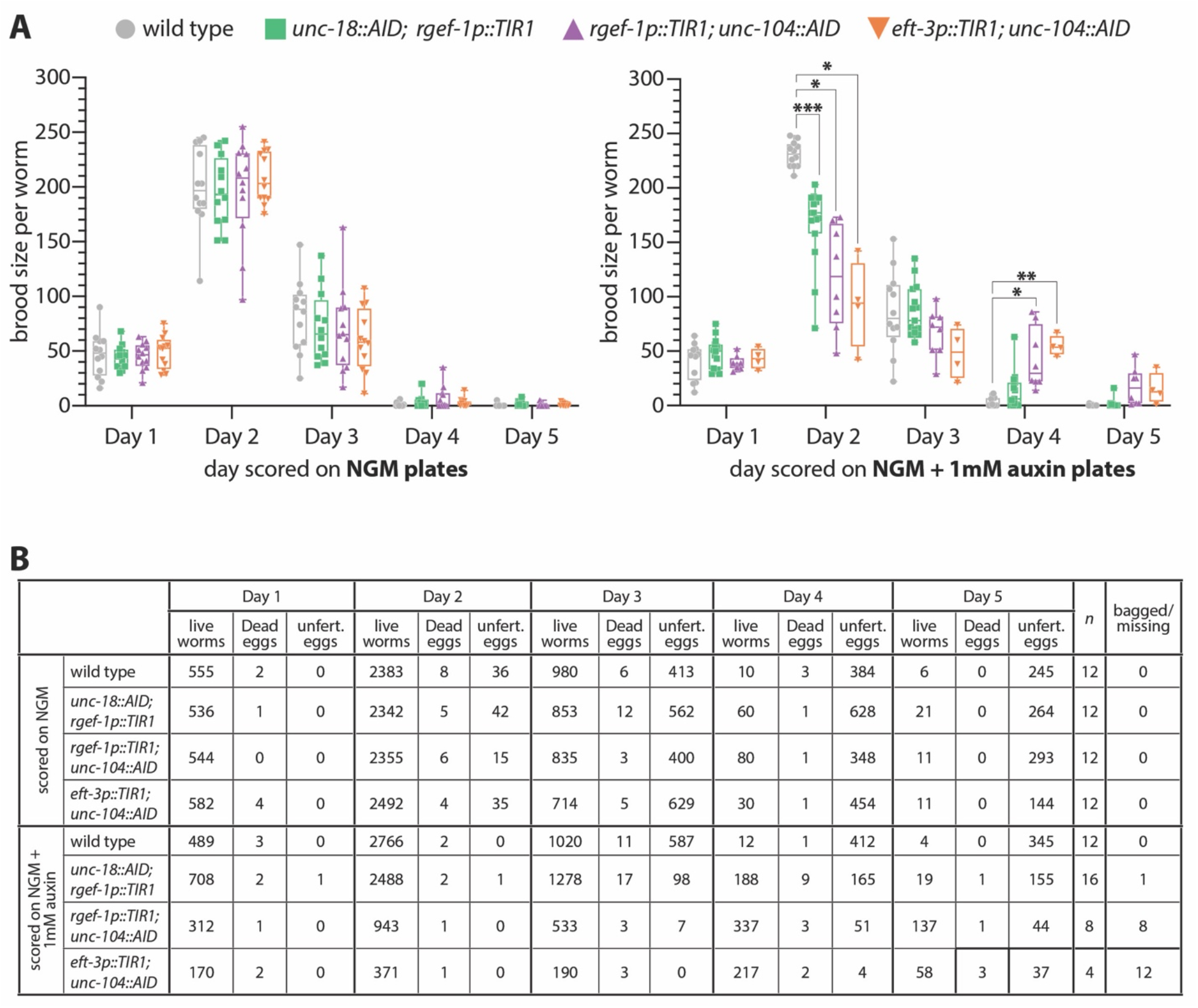
Auxin-dependent depletion of *unc-18* does not significantly alter fertility. **(A)** Brood size calculations and **(B)** progeny counts of lives worms, dead eggs and unfertilized (unfert.) eggs for wild type, *unc-18::AID; rgef-1p::TIR1, rgef-1p::TIR1; unc-104::AID*, and *eft-3p::TIR1; unc-104::AID* on nematode growth media (NGM) with and without 1mM auxin. N value indicates number of parental hermaphrodites scored and bagged/missing indicates the number of worms that bagged or disappeared from the plate during the 5 days of scoring. * indicates P<0.01, ** indicates P<0.001, *** indicates P<0.0001, Dunnett’s multiple comparisons test.

In the absence of auxin, all genotypes displayed brood sizes that were indistinguishable from wild type worms through all five days of scoring (Figure 2A). In particular, wild type worms display similar brood sizes both on and off auxin throughout all five days. Additionally, the number of dead eggs was also similar between the on and off auxin wild type worms suggesting that oocyte viability is not being altered (Figure 2B). Further, the cumulative sum of the brood size for wild type both on and off auxin (no auxin average cumulative brood size for 12 worms: 329.417 ± 62.2 SD; auxin average cumulative brood size for 12 worms: 360.917 ± 25.6 SD) is comparable to published results showing that wild type animals have a cumulative total brood size of ∼300 worms (Ward and Carrel 1979; Muschiol *et al*. 2009). Moreover, previous studies have found as well that auxin exposure does not alter the viability or fertility of wild type worms (Zhang *et al*. 2015; Martinez *et al*. 2020; Ashley *et al*. 2021). Taken together, this mounting evidence supports a conclusion the auxin does not affect the fertility or germline progression in wild type worms.

In the presence of auxin, *unc-104::AID* worms displayed slight changes in fertility (Figure 2A). The progeny laid on auxin plates during the first 24 hours (Day 1) displayed no significant changes in brood size for any of the genotypes examined compared to wild type animals. Over the next 24 hours (Day 2) on auxin, *unc-104::AID* displayed significant reductions in brood size compared to wild type (*rgef-1p::TIR1; unc-104::AID* P=0.0135; *eft-3p::TIR1; unc-104::AID* P=0.0158, Dunnett’s multiple comparisons). In contrast, on days four and five *unc-104::AID* displayed larger brood sizes than wild type animals with these changes being only significantly different on Day 4 (Day 4: *rgef-1p::TIR1; unc-104::AID* P=0.0447; *eft-3p::TIR1; unc-104::AID* P=0.0036, Dunnett’s multiple comparisons; Day 5: *rgef-1p::TIR1; unc-104::AID* P=0.1344; *eft-3p::TIR1; unc-104::AID* P=0.1944, Dunnett’s multiple comparisons). This result suggests that depletion of UNC-104 might cause a slight delay in oocyte progression through the germline allowing more oocytes to be laid later during the hermaphrodite reproductive lifespan. Further experiments are necessary to determine if indeed this is a delay or if something else is contributing to the discrepancies in brood size compared to wild type animals.

In addition to having significant changes in brood size on auxin, *unc-104::AID* worms also displayed a high number of worms with a bag of worms phenotype, where worms retained their progeny within their body cavity, and worms that went missing over the five days of scoring (Figure 2B). Additionally, *unc-104::AID* worms on auxin displayed behavioral defects where instead of staying within the bacteria lawn on the plate these worms exhibited a wandering pattern outside of the bacterial lawn that was abnormal for hermaphrodites. Typically, hermaphrodite worms remain within the bacteria lawn and rarely wander to the edges of the plate (Lipton *et al*. 2004). However, the *unc-104::AID* worms on auxin would drag themselves in non-sinusoidal patterns to the edges of the plates where they would stick to the plastic and desiccate before dying. Taken together, this data suggests that the *unc-104::AID* worms may have both a behavioral defect and a slight fertility defect or delay. Due to these defects, we decided to exclude *unc-104::AID* for use any future live imaging experiments and processed with analysis on *unc-18::AID* worms.

### UNC-18::AID depletion has minimal effects the hermaphrodite germline

In the presence of auxin, *unc-18::AID* worms displayed slight changes in brood size, but the brood viability (dead eggs) remains indistinguishable from wild type (Figure 2). The progeny laid from *unc-18::AID* worms during the first 24 hours (Day 1) on auxin exhibited no significant changes in brood size. In contrast, the progeny on auxin at Day 2 displayed significant reductions in brood size compared to wild type (*unc-18::AID; rgef-1p::TIR1* P=0.0002, Dunnett’s multiple comparisons). Notably, this decrease in brood size on Day 2 did not correlate with an increase in the number of dead eggs, indicating that oocyte viability is not the likely cause of this brood size reduction (Figure 2B). Further, over the next three days of scoring, the brood size of *unc-18::AID* worms was indistinguishable from wild type worms on auxin suggesting that this slight reduction only effects the 24-48hr progeny. Additionally, *unc-18::AID* worms displayed none of the behavioral or worm bagging defects that were seen with *unc-104::AID* animals (Figure 2B).

The slight reduction in the 24-48hr brood when UNC-18 is depleted could be explained by changes in germline progression. To assess for changes in germline progression, we used an EdU pulse-chase experiment where worms were soaked in EdU (“pulse”) to allow EdU incorporation into nuclei undergoing DNA replication, including those in the pre-meiotic tip. Then, worms were dissected at 0, 10, 24, and 48 hours post EdU soaking to “chase” the EdU staining in the germline (see Methods) (Crittenden *et al*. 2006; Jaramillo-Lambert *et al*. 2007; Morgan *et al*. 2010; Fox *et al*. 2011; ROSU *et al*. 2011; Seidel and Kimble 2015; Almanzar *et al*. 2021). To score the EdU progression, the *C. elegans* germline was divided up into four different regions based on nuclear DNA morphology: pre-meiotic to transition zone (PMT-TZ), early pachytene to mid pachytene (EP-MP), mid pachytene to late pachytene (MP-LP), and diakinesis to germline end (diakinesis+) (Hillers *et al*. 2017). Since the results of our fertility assays (Figure 2) and those of multiple studies have found that auxin has no effect on the viability and/or fertility in wild type worms (Zhang *et al*. 2015; Martinez *et al*. 2020; Ashley *et al*. 2021), we performed all the EdU experiments comparing wild type worms in the absence of auxin to *unc-18::AID* worms in presence of auxin, unless otherwise indicated. For our EdU pulse-chase experiments, both wild type and *unc-18::AID* displayed very similar patterns of EdU staining throughout the time course with nuclei appearing to progress at similar rates (Figure 3A). Germlines initially show staining in the PMT-TZ region, which is indicative of the mitotic and meiosis S phase DNA replication occurring in this region. Then, the EdU front slowly progresses through each germline stage from EP-MP at 10 hours, MP-LP at 24 hours and past diakinesis (Diakinesis+) at 48 hours. Overall, the progression of nuclei through the germline is not grossly altered by auxin mediated depletion of UNC-18::AID.

**Figure 3.**
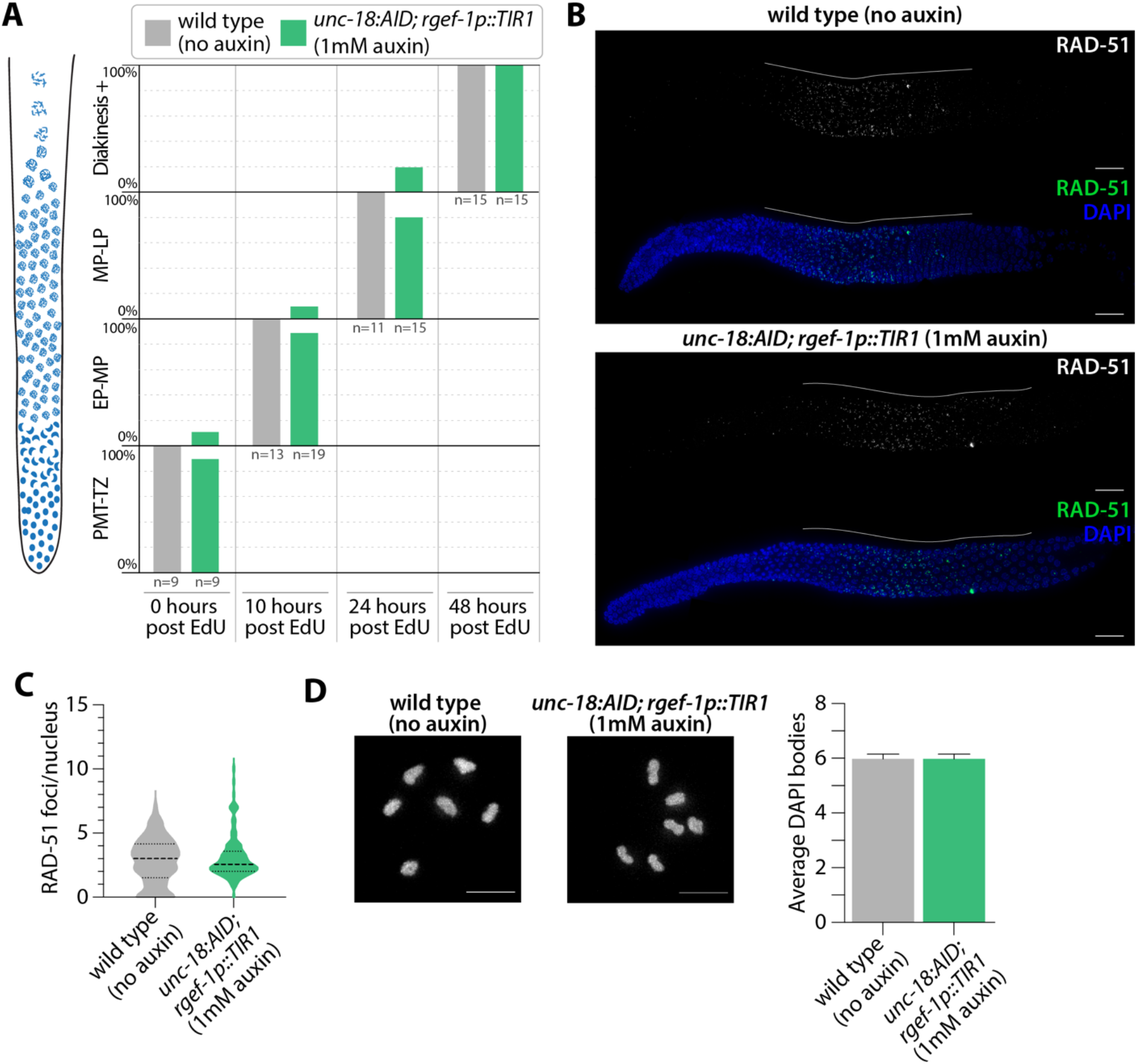
Hermaphrodite germline progression and meiotic crossover formation are unaffected by auxin-depletion of *unc-18*. **(A)** Quantification of the nuclear progression through the germline in wild type with no auxin exposure and *unc-18::AID; rgef-1p::TIR1* on 1mM auxin plates. Each gonad was scored based on the germline position of the last EdU stained nucleus from gonads dissected at 0, 10, 24, and 48 hours post EdU labeling. For this analysis the germline (diagrammed on the left) was divided into four regions: premeiotic tip to transition zone (PMT-TZ), early pachytene to mid pachytene (EP-MP), mid pachytene to late pachytene (MP-LP), and diakinesis to end of the germline (Diakinesis+). N values for number of worms scored are displayed on the plot below each bar. **(B)** Representative images of dissected gonads from wild type (no auxin exposure) and *unc-18::AID; rgef-1p::TIR1* (1mM auxin exposure) hermaphrodites stained for RAD-51 (green) and DNA (DAPI, blue). Scale bar represents 20µm and white line indicates the length of the RAD-51 staining within the germline. **(C)** Quantification of the number of RAD-51 foci per nucleus for wild type (no auxin exposure, n=8 gonads, 438 nuclei) and *unc-18::AID; rgef-1p::TIR1* (1mM auxin exposure, n=9 gonads, 608 nuclei). **(D)** Representative image and quantification of diakinesis chromosomes (DAPI staining bodies) from wild type (no auxin exposure, n=30 oocytes) and *unc-18::AID; rgef-1p::TIR1* (1mM auxin exposure, n=30 oocytes).

To determine if meiotic recombination is altered by depletion of UNC-18::AID, we assayed for the initiation of recombination through the quantification of meiotic double-strand DNA breaks (DSBs) using the recombinase protein RAD-51. At the onset of meiosis, the topoisomerase-like enzyme SPO-11 induces DSBs from which a subset are repaired via homologous recombination into crossover events, which ensure accurate segregation of the homologs during meiosis I (reviewed in Hunter 2015). In *C. elegans*, DSBs are formed during the transition zone through early pachytene, and then RAD-51 is loaded on to the resected single-stranded DNA ends of the DSB site (Hillers *et al*. 2017). As the nuclei progress through pachytene, the DSB is repaired by homologous recombination and RAD-51 is removed from the DSB site (Hillers *et al*. 2017). Using immunofluorescence, we quantified the amount of RAD-51 foci within the pachytene region of the germline in both wild type and *unc-18::AID* worms (Figure 3B,C). *unc-18::AID* worms did not display any significant difference in RAD-51 foci compared to wild type (P=0.7072, Mann-Whitney). Thus, formation of DSBs and subsequent off-loading of RAD-51 is unaffected by depletion of *unc-18::AID* by auxin.

To determine if UNC-18::AID depletion had any effect an crossover formation we assayed for the presence of crossover events by quantifying the number of DAPI staining DNA bodies at diakinesis. Since *C. elegans* have 6 pairs of homologous chromosomes which are connected by crossover events to form bivalents at the diakinesis stage of meiotic prophase I, greater than 6 DAPI staining DNA bodies in diakinesis nuclei indicates errors in meiotic prophase I (*e*.*g*. lack of crossovers or chromosome fragmentation from unrepaired DSBs). Both wild type and *unc-18::AID* worms had an average of 6 DAPI staining DNA bodies at diakinesis (P>0.99, Mann-Whitney), thereby indicating depletion of UNC-18::AID has minimal effects on meiotic recombination. Taken together, subtle differences in both germline progression and brood size in UNC-18::AID depletion do not have obvious negative consequences on meiosis.

### Conditional immobilization of hermaphrodites for live imaging

To immobilize worms for live imaging, 5-6 adult hermaphrodites carrying a fluorescent reporter and *unc-18::AID; rgef-1p::TIR1* were selected from 1mM auxin plates. These worms were placed into a 1-2µL drop of imaging M9 media containing 25mM serotonin and 1mM auxin on a poly-lysine coated coverslip. Previous studies have shown that serotonin is important to maintain motion within the germline during live imaging (Rog and dernburg 2013; Rog and Dernburg 2015; Pattabiraman *et al*. 2017). Further, the addition of auxin within this media ensures a continued depletion of UNC-18::AID throughout the imaging. Using M9 media, we made 7-9% agarose pads and this pad was then transferred and carefully laid over the top of the worm. Whatman paper was used to wick away any excess liquid between the coverslip and agarose pad. Then, the coverslip-worm-agar pad sandwich was sealed to a slide using Vaseline to prevent drying out of the agarose pad during live imaging (Figure S1).

For brightfield imaging, worms were imaged every 90 seconds for a total of 60 minutes. During that time, mounted hermaphrodite worms are able to subtly move their heads and some of the worms continued to ovulate oocytes that would stack up in a pile next to the worm (2-3 oocytes/60 min, 7 worms, Figure 4A, Movie S1). This continued ovulation of the hermaphrodite is an excellent indicator that germline progression is not being impeded by having the worms mounted underneath an agar pad. Additionally, the movement of the germline can be seen by directly looking at the germline nuclei using a fluorescently tagged component of the synaptonemal complex (SC), *SYP-2::GFP*, which is a meiotic chromosome structure that assembles between homologous chromosomes in the germline from late transition zone to diakinesis (Figure 4B, Movie S2). To minimize motion of the worm at 60x, we included 0.08% tricaine and 0.008% tetramisole anesthetics, which is at a concentration nearly 10-fold lower than previous studies (Wynne *et al*. 2012; Rog and Dernburg 2015; Pattabiraman *et al*. 2017; Rog *et al*. 2017). In comparison to the previous brightfield timelapse image without anesthetics (Movie S1), we found that combining this very small amount of anesthetics with knockdown of UNC-18::AID removed all of the residual head motion (Movie S2, S3). Further, this combination of low concentration anesthetics paired with UNC-18::AID knockdown allowed for stable imaging of SYP-2::GFP worms every 5 minutes for up to 2 hours and did not interfere with the post-imaging recovery of the worms. Using SYP-2::GFP, we observed the previously described motion of the SC within each germ cell nucleus of living worms as well as the proximal movement of the nuclei away from the distal tip cell (Movie S2, S3) (Wynne *et al*. 2012; Rog and Dernburg 2013; Rog and Dernburg 2015; Pattabiraman *et al*. 2017). Notably, we can also observe the progressive motion of the diakinesis oocytes, which further indicates that oocyte progression is unimpeded by this immobilization method (Movie S3).

**Figure 4.**
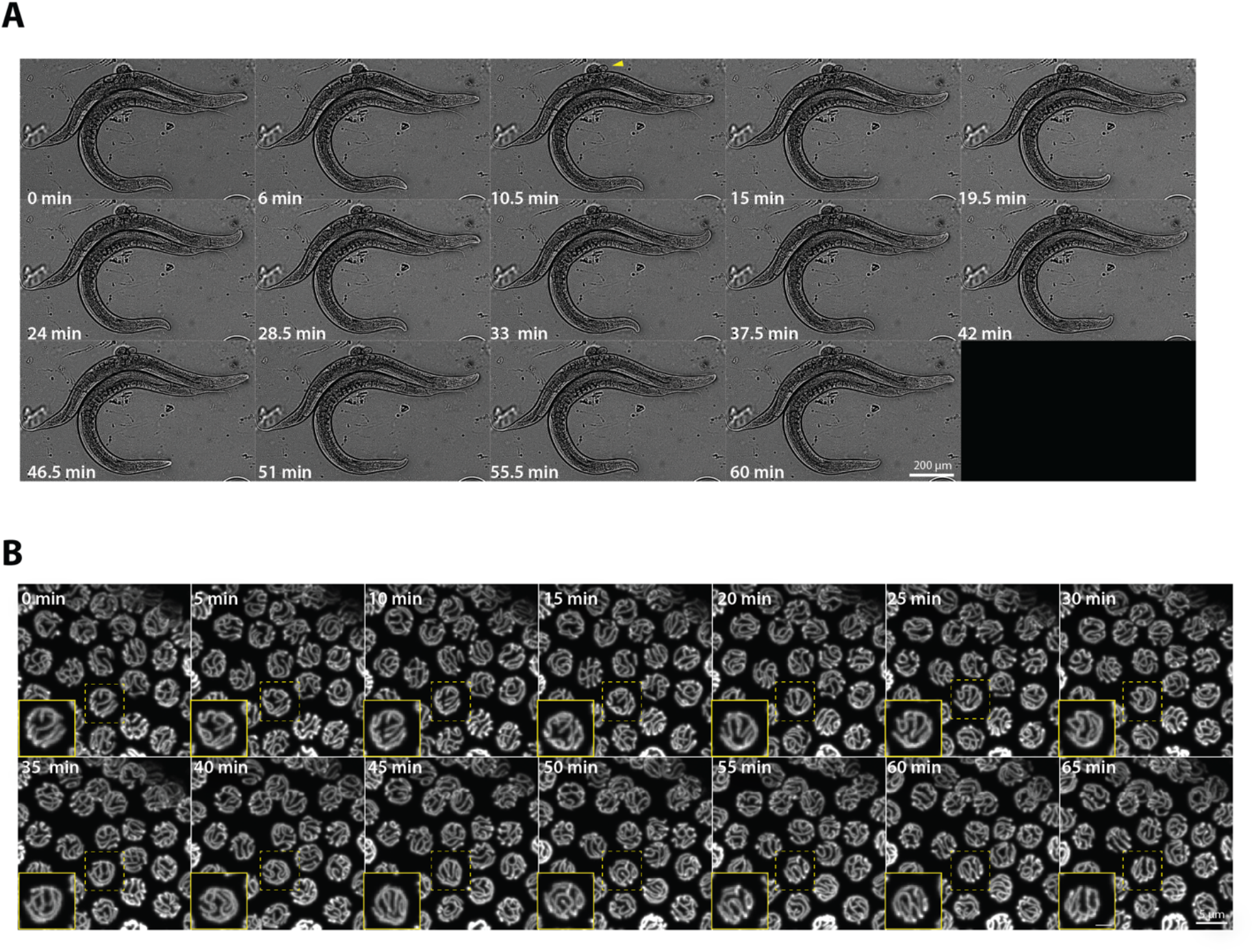
Conditional immobilization of hermaphrodites for live imaging. **(A)** Brightfield timelapse montage of an immobilized hermaphrodite worm at 40x magnification with images captured every 90 seconds for 60 minutes. The montage displays every third frame of the timelapse. Yellow arrowhead indicates ovulation of an egg by the immobilized hermaphrodite. **(B)** SYP-2::GFP timelapse montage of the hermaphrodite germline at 60x magnification with images captured every 5 minutes for 65 minutes. The mid-pachytene region of the germline is shown and germline is moving from left to right in each image. The complete movies can be viewed in Movies S1 and S2. The yellow dashed box indicates the nucleus that is enlarged in the inset panel (yellow outline) with the scale bar in the inset panel representing 2 µm

### unc-18::AID reversibly immobilizes male worms for live imaging

One of the unique features of our conditional immobilization system using *unc-18::AID; gref-1p::TIR1* is that it works well with whole, intact male worms. From our immobilization experiments, we found that male worms require a higher concentration of auxin to induce a more robust immobilization than hermaphrodite worms (Figure 5A). Male worms grown on plates containing 10mM auxin exhibited a more severe immobilization phenotype and a tighter distribution of average speeds than male worms on 1mM auxin (median average speed 1.251 and 4.809 pixels/second, respectively). Further, male worms removed from auxin for 18-24 hours recovered sinusoidal movements close to wild type levels regardless of the initial auxin concentration. These results suggest a sexual dimorphic difference in auxin sensitivity in *C. elegans*, where male worms may absorb and process auxin differently than hermaphrodite worms. Future studies focused on auxin processing in both *C. elegans* sexes may reveal the mechanisms behind this intriguing sexual dimorphism. Based on these analyses which revealed a more robust immobilization at higher auxin concentrations, we proceeded with using 10mM auxin to immobilize male worms for live imaging.

**Figure 5.**
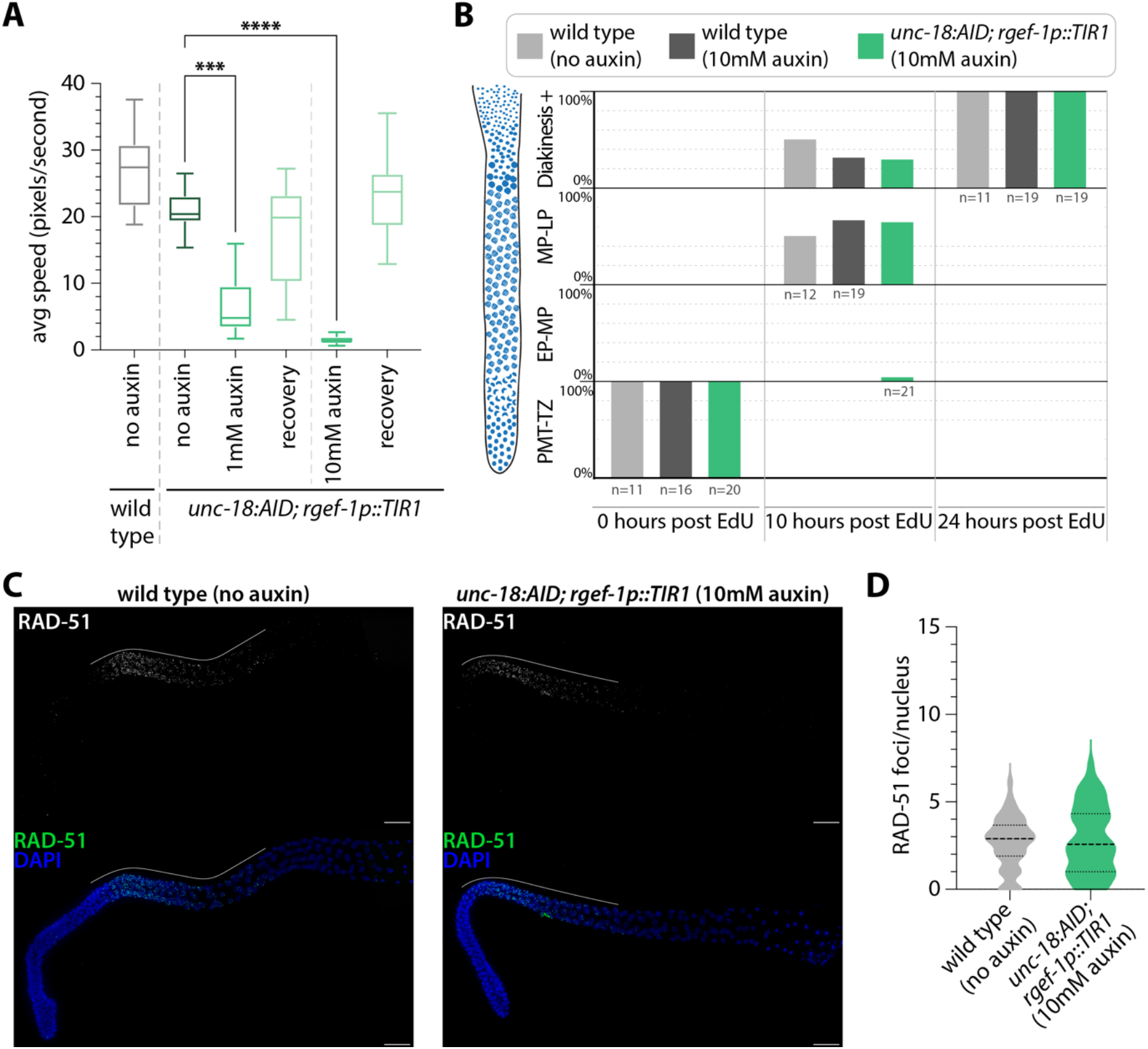
Conditional immobilization of males following auxin dependent depletion of *unc-18*. **(A)** Quantification of the average speed in pixels per second of wild type (n= 24) and *unc-18::AID; rgef-1p::TIR1* on plates containing no auxin (n=29), 1mM auxin (n=25), recovery from 1mM auxin (n=21), 10mM auxin (n=30), and recovery from 10mM auxin (n=21). The recovery category indicates worms that have been off auxin for 18-24 hours prior to tracking worm motion. *** indicates P<0.0001 and **** indicates P<0.00001, Kruskal-Wallis. **(B)** Quantification of the nuclear progression through the germline in wild type with no auxin exposure and on 10mM auxin plates and *unc-18::AID; rgef-1p::TIR1* on 10mM auxin plates. Each gonad was scored based on the germline position of the last EdU stained nucleus from gonads dissected at 0, 10 and 24 hours post EdU labeling. For this analysis, the germline (diagrammed on the left) was divided into four regions: premeiotic tip to transition zone (PMT-TZ), early pachytene to mid pachytene (EP-MP), mid pachytene to late pachytene (MP-LP), and diakinesis to end of the germline (Diakinesis+). N values for number of worms scored are displayed on the plot below each bar. **(C)** Representative images of dissected gonads from wild type (no auxin exposure) and *unc-18::AID; rgef-1p::TIR1* (1mM auxin exposure) males stained for RAD-51 (green) and DNA (DAPI, blue). Scale bar represents 20µm and white line indicates the length of the RAD-51 staining within the germline. **(D)** Quantification of the number of RAD-51 foci per nucleus for wild type (no auxin exposure, n=8 gonads, 104 nuclei) and *unc-18::AID; rgef-1p::TIR1* (10mM auxin exposure, n=8 gonads, 94 nuclei).

Similar to hermaphrodites, we wanted to ensure that depleting UNC-18::AID with the neuron expressed *TIR1* (*rgef-1p::TIR1*) did not cause detrimental changes to the *C. elegans* germline. To determine if depletion of UNC-18::AID altered the progression of nuclei in the male germline, we used EdU labeling to track germ cell nuclei progression. Previous studies have demonstrated that the male germline progresses much faster than the hermaphrodite germline (24 hours to complete spermatogenesis versus 48-72 hours to complete oogenesis) (Jaramillo-Lambert *et al*. 2007). To monitor EdU-labeled nuclei progression within the germline, we dissected the germlines out of male worms following EdU soaking at 0, 10, and 24 hours. Wild type male worms both in the presence and absence of 10mM auxin displayed similar rates of germline progression, suggesting that auxin does not interfere with germline progression in males (Figure 5B). In addition, depletion of UNC-18::AID did not appear to cause gross changes in male germline progression. Thus, overall progression of the male germline is relatively unaltered by the presence of both 10mM auxin and depletion of UNC-18::AID.

In addition to normal germline progression, the initiation and repair of DSBs appear unaffected by UNC-18 depletion in the male germline. Analysis of RAD-51 foci using immunofluorescence revealed no significant changes in the number of RAD-51 foci per nucleus during pachytene (Figure 5C). Wild type worms in the absence of auxin had an average of 2.66 RAD-51 foci per nucleus (± 1.5 SD) and *unc-18::AID; rgef-1p::TIR1* worms on auxin have an average of 2.79 RAD-51 foci per nucleus (± 1.9 SD). Further, RAD-51 foci appeared to be off-loaded from the nuclei such that foci did not persist into the later stages of spermatogenesis (Figure 5C). Taken together, DSB formation and repair does not appear to exhibit any defects in the presence of 10mM auxin and depletion of UNC-18::AID.

To enable live imaging of male worms, we mounted the worms using the same steps as described above for the hermaphrodite worms except we used 10mM auxin in the M9 media instead of 1mM auxin to maintain depletion of UNC-18::AID during imaging (Figure S1). All live imaging experiments were performed using the same imaging settings as the hermaphrodites for except for the brightfield timelapses, which due to the smaller size of the male worms were captured at 20x magnification instead of 10x (Figure 6). Notably, the same mounting and immobilization method worked as efficiently to immobilize the male worms as it did with the hermaphrodite worms (Figure 6A, Movie S4). Further, using SYP-2::GFP with the *unc-18::AID; rgef-1::TIR1* conditional immobilization technique, we were able to observe both the directional motion of germline progression and the chromosome motion within each nucleus (Figure 6B, Movie S5, S6).

**Figure 6.**
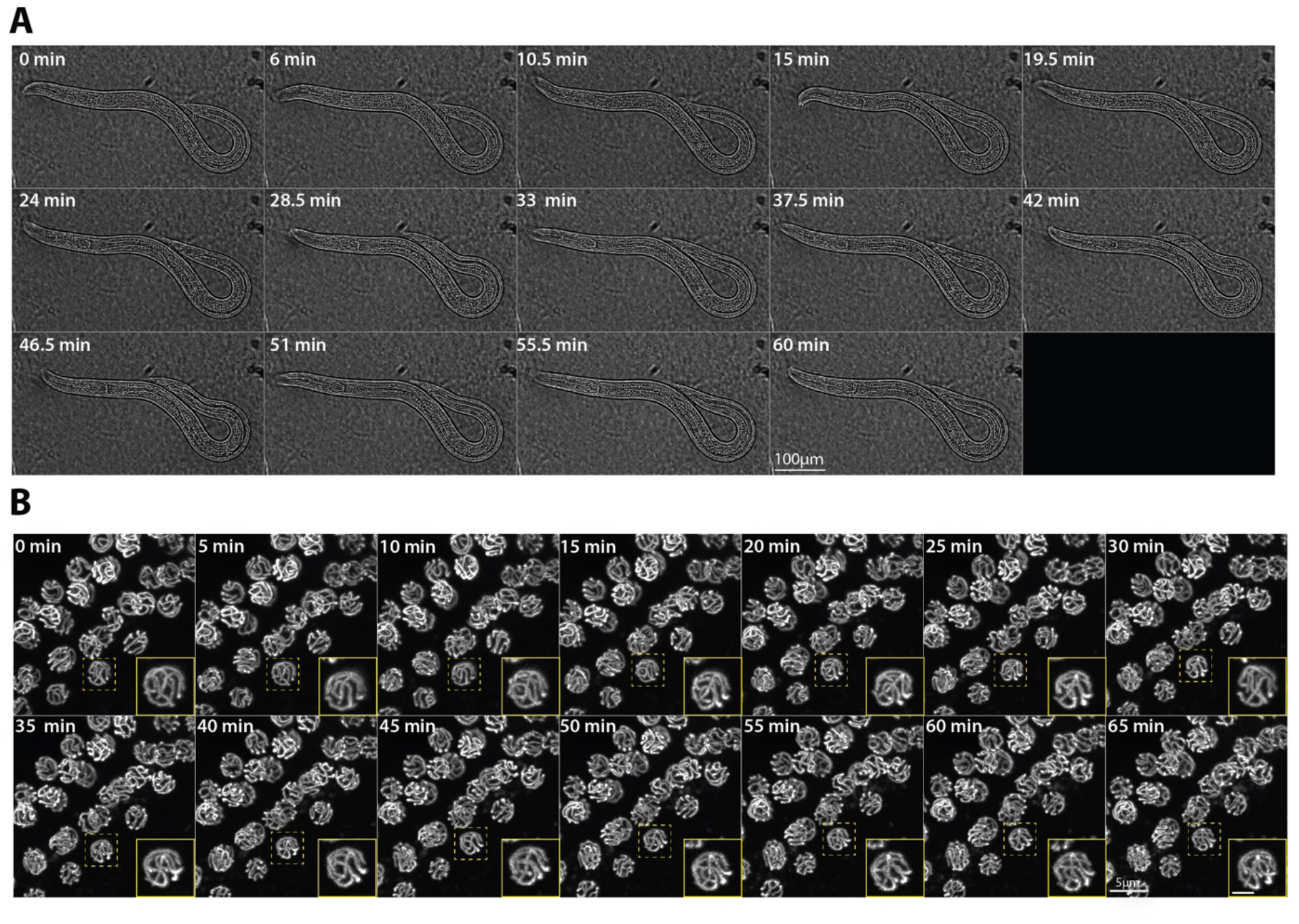
Conditional immobilization of males for live imaging. **(A)** Brightfield timelapse montage of an immobilized male worm at 20x magnification with images captured every 90 seconds for 60 minutes. The montage displays every third frame of the timelapse. **(B)** SYP-2::GFP timelapse montage of the male germline at 60x magnification with images captured every 5 minutes for 65 minutes. The mid-pachytene region of the germline is shown and germline is moving from bottom left to top right in each image. The complete movies can be viewed in Movies S4 and S5. The yellow dashed box indicates the nucleus that is enlarged in the inset panel (yellow outline) with the scale bar in the inset panel representing 2 µm.

## DISCUSSION

The transparent nature of the *C. elegans* worm makes this model organism ideal for live imaging studies, however, effectively and reliably immobilizing the worm without injury has been a challenge for many *C. elegans* labs seeking to do live imaging experiments. We developed and validated a new tool that enables conditional immobilization of *C. elegans* for live imaging the germline. This conditional immobilization tool uses the auxin inducible degron system, which we show works for immobilizing both hermaphrodite and male worms. Notably, we found that depletion of the gene product responsible for this immobilization phenotype does not cause any significant changes within the germline of either sex. Finally, with this tool we were able to demonstrate that both male and hermaphrodite worms can be minimally restrained as whole animals with an agar pad and imaged live for at least two hours (Movies S4 and S6).

The conditional immobilization technique described here to immobilize worms enhances the existing toolkit for live imaging worms. While many modalities exist from microfluidic chips to pharmaceuticals for immobilization of the worm, here we present an accessible genetic tool that can be used to easily immobilize worms and can be implemented in any lab without needing to purchase specialized equipment or use hazardous chemicals. Notably, the use of anesthetics has been widespread for live imaging studies; however, male *C. elegans* are known to respond differently than hermaphrodites to different chemicals and toxins including the widely used anesthetic Levamisole (Lopes *et al*. 2008; Ruszkiewicz *et al*. 2019). Thus, methods used to immobilize hermaphrodite worms may not be as effective against male worms and have inhibited live imaging of sexual dimorphisms and male-specific processes. A recent study demonstrated that male worms can be immobilized for live imaging of the spermatocyte divisions using an agar pad approach (Fabig *et al*. 2020). After incorporating a modified version of this agar pad approach with our conditional immobilization tool (see Methods), we were able to consistently and reliably mount male worms without any of the technical challenges associated with the agar pad handling or placement. Specifically with our combination of these techniques, we found that regardless of how the agar pad was placed over the males, all of the mounted male worms were suitable for imaging and remained immobile. Overall, multiple options for immobilization of male worms increases the possibility of live imaging experiments that can be performed in a multitude of laboratory settings with different resources.

Our conditional immobilization tool works to efficiently immobilize both males and hermaphrodites for live imaging worms for sex-specific comparison experiments. For the *C. elegans* germline, an increasing number of studies are indicating hermaphrodites and males have multiple sexually dimorphic features that lead to differential regulation of genome integrity germline (Jaramillo-Lambert *et al*. 2007; Cahoon and Libuda 2019; Kurhanewicz *et al*. 2020; Li *et al*. 2020). To our knowledge, the live images of the SC in males may represent the first time the SC has been observed within living *C. elegans* spermatocytes. This conditional immobilization tool enables future experiments examining the dynamics of the SC between the sexes and has the potential to uncover new sexual dimorphic features of the *C. elegans* germline. Further, this method can be used to examine additional sexual dimorphisms within the germline such as germline progression, chromosome compaction, meiotic cell divisions and the endogenous RNAi system. Moreover, sexual dimorphisms occur outside of the germline and this system can be also used to live image the development of other tissues within the worm such as the digestive tract and muscles, both of which display sexually dimorphic features during development and within the adult animal (Emmons 2014).

Our immobilization system has both flexibility for use in a variety of experiments and the ability for future modifications and expansions. To enable live imaging of any existing tagged fluorescence reporter strains, the components of our live imaging system can easily be incorporated through either genetic crosses or CRISPR-Cas9 injections (see Methods and Table S1 for CRISPR details). Moreover, there are likely more genes within the *C. elegans* genome that could be tagged with AID and used to induce analogous immobilization to UNC-18 depletion. Future expansion of the tool kit for the conditional immobilization of worms will provide even greater genetic flexibility for use of this system in a multitude of live imaging studies.

## ACKNOWLEDGEMENTS

We thank the CGC for strains, which is funded by NIH Office of Research Infrastructure Programs (P40 OD010440). We thank members of the Libuda Lab, especially N. Kurhanewicz, A. Naftaly and E. Toraason, for discussion and comments on the manuscript. This work was supported by the National Institutes of Health R35GM128890 to DEL and a Jane Coffin Childs Postdoctoral Fellowship to CKC. DEL is also a Searle Scholar and recipient of a March of Dimes Basil O’Connor Starter Scholar award.

---

**Figure S1.**
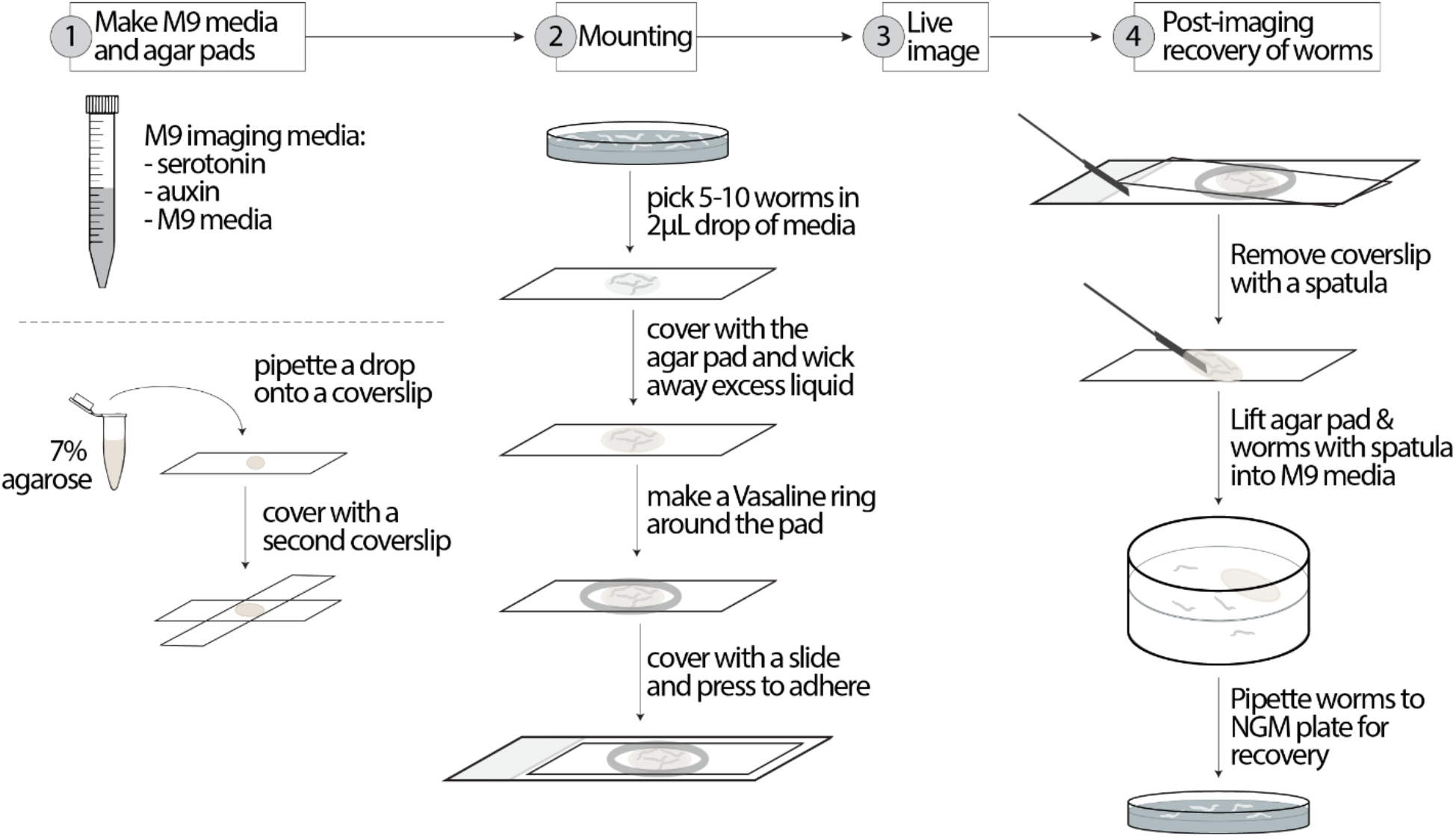
Diagram of live imaging mounting and post-imaging recovery. For a comprehensive description see the Methods section “*Conditional immobilization for live imaging”*.

**Movie S1. Brightfield 10x timelapse movie of an immobilized hermaphrodite worm**. A hermaphrodite worm containing *unc-18::AID; rgef-1p::TIR1* that was immobilized on 1mM auxin and mounted under an agar pad. Images were captured every 90 seconds for 60 minutes.

**Movie S2. SYP-2::GFP in an immobilized hermaphrodite worm**. Timelapse images were taken at 60x and images were captured every 5 seconds for 65 minutes. Movie was stabilized for viewing (see Methods for details).

**Movie S3. Whole germline view of SYP-2::GFP in an immobilized hermaphrodite worm**. Timelapse images were taken at 60x and images were captured every 5 seconds for 2 hours. The start of the germline occurs out of view of the camera on the left and germline progresses from left (transition zone/early pachytene) to right (late pachytene/diplotene) within the movie. Then, the germline makes a U-turn bend, which is out of view of the camera. The germline comes back into view indicated by the arrowhead showing the appearance of the diakinesis oocytes on the bottom right of the movie, which are moving from right to left. Movie was stabilized for viewing (see Methods for details).

**Movie S4. Brightfield 20x timelapse movie of an immobilized male worm**. A male worm containing *unc-18::AID; rgef-1p::TIR1* that was immobilized on 10mM auxin and mounted under an agar pad. Images were captured every 90 seconds for 60 minutes.

**Movie S5. SYP-2::GFP in an immobilized male worm**. Timelapse images were taken at 60x and images were captured every 5 seconds for 65 minutes. Movie was stabilized and corrected for photobleaching for viewing (see Methods for details).

**Movie S6. Whole germline view of SYP-2::GFP in an immobilized male worm**. Timelapse images were taken at 60x and images were captured every 5 seconds for 2 hours. Arrowhead indicates the approximate start of the germline within the mid-bottom region of the movie. Germline progression initially starts off moving to the left, then the germline makes a U-turn bend at the transition zone on the left side of the movie. After this bend, the germline progression moves from the left (early pachytene) to right (late pachytene/diplotene) within the movie. The end of the germline (condensation and mature sperm) occurs farther off to the right outside of the camera view at this magnification. Movie was stabilized and corrected for photobleaching for viewing (see Methods for details).

**Table S1:**
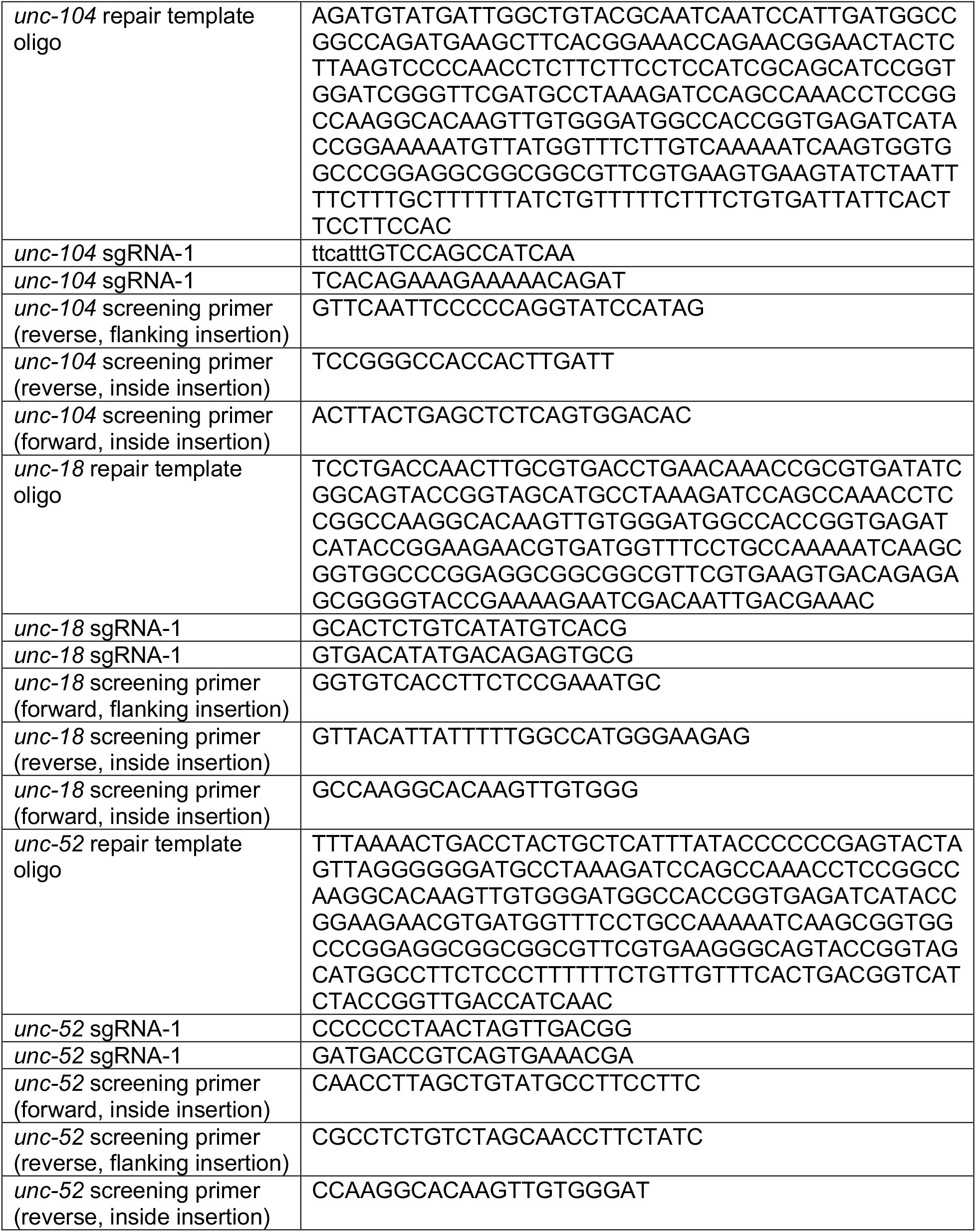
Primer sequences used for CRISPR/Cas9 tagging *unc-104, unc-18*, and *unc-52* with auxin inducible degron tag ^*^.All DNA sequences are presented in the 5′ to 3′ direction

